# Individual differences in perceptual capacity depend on aperiodic slope, not alpha oscillations

**DOI:** 10.1101/2025.07.21.666053

**Authors:** Jessica A. Elliott, Jason B. Mattingley, Joshua O. Eayrs, Anthony M. Harris

## Abstract

Visual processing is subject to strict capacity limitations that restrict the number of stimuli that can be individuated, that is, distinguished as separate objects, to around 3-4 items at a time. The neural mechanisms that give rise to this ‘subitizing’ limit are unclear. Here, we tested an account of the subitizing limit that attributes it to the amount of information that can be processed within one cycle of the 8-14 Hz alpha oscillation. This account proposes that the subitizing limit should be correlated with an individual’s alpha oscillation frequency. We pitted this account against one based on neural excitatory-inhibitory balance, as indexed by the slope of the aperiodic component of the electrophysiological power spectrum. To test these accounts, we had human participants (N=51; 37 females) complete a visual enumeration task while we measured their brain activity with electroencephalography (EEG). We extracted aperiodic and alpha-band activity from the pre-stimulus period and correlated these with model-based estimates of individuals’ subitizing capacity. Alpha oscillations were not correlated with subitizing capacity within-participants or as individual differences, but they were associated with other elements of enumeration performance, including baseline reaction times and the slope of the performance decrement outside the subitizing range. By contrast, individual differences in aperiodic slope reliably predicted individuals’ subitizing capacity. These results suggest that differences in neural excitatory:inhibitory balance are the source of individuation-related capacity limits, not alpha-frequency sampling.

**Significance Statement:** Busy, multi-element visual scenes are common in daily life. Human visual perception, however, is capacity limited, individuating only 3-4 objects at a time. The source of these capacity limitations is currently unknown. This work examines the neural correlates of perceptual capacity within and between individuals, comparing a rhythmic ‘pulsed inhibition’ account with one based on the arhythmic balance of neural excitation and inhibition. Our results demonstrate that individuals with more arhythmic inhibition demonstrate greater perceptual capacity. These results help to explain how excitation and inhibition across populations of neurons sculpts the capacity of the neural system and suggest targets for future interventions to improve perceptual capacity with arhythmic brain stimulation.

---

The phase, amplitude and frequency of neural oscillations all predict the processing quality of incoming sensory signals (VanRullen, 2016; Clayton et al., 2018). Non-oscillating, aperiodic, neural states also predict sensory processing (Cunningham et al., 2023; Deodato and Melcher, 2023; Williams et al., 2024). Little is known, however, about how these indicators of electrophysiological neural state relate to inherent capacity limitations in human vision. One such limit, the subitizing capacity (Jevons, 1871; Kaufman et al., 1949), reflects a constraint on the number of objects that can be individuated (recognised as distinct from their surroundings) at any one time (Xu and Chun, 2009). The subitizing limit is demonstrated in participants’ ability to enumerate 3-4 items with uniform efficiency, with performance degrading steadily beyond this range. The 3-4 item subitizing limit has been described as a measure of ‘perceptual capacity’ as it shares mechanisms with other visual capacity limits, such as those observed in multi-object tracking and change blindness (Eayrs and Lavie, 2018, 2021). Here, we characterised how individual differences in subitizing limit relate to pre-stimulus 8-14 Hz alpha oscillations and aperiodic neural activity.

The influence of pre-stimulus alpha oscillations on visual processing is well established. Alpha oscillations modulate visual processing in a rhythmic manner (Busch et al., 2009; Mathewson et al., 2009; Harris et al., 2018; Williams et al., 2024). Many experiments have shown that the peak frequency of alpha oscillations is associated with the temporal resolution of visual processing (Samaha and Postle, 2015; Gulbinaite et al., 2017; Ronconi et al., 2022; Samaha and Romei, 2023). Results such as these support the hypothesis that perceptual ‘sampling’ is instantiated by alpha oscillations. According to this idea, alpha oscillations cycle sensory systems between active and inhibited states, and all sensory signals arriving during an active state are perceived as one ‘perceptual snapshot’. This idea of discrete perception can be thought of through the analogy of discrete frames in a movie that give rise to the impression of continuous motion (VanRullen and Koch, 2003; Lundqvist and Wutz, 2022).

It has been suggested that capacity limits on object individuation may emerge from constraints on the amount of information that can be processed within one cycle of the alpha wave (Wutz and Melcher, 2014). This theory proposes that visual information is processed in a serial-like manner (Wutz et al., 2012; Wutz and Melcher, 2013), with 3-4 items being processed within one alpha cycle. To date, however, no study has tested the hypothesis resulting from this claim, that individual differences in subitizing should be predicted by an individual’s specific alpha oscillation frequency. Such a relationship would provide strong support for an alpha-based model of perceptual capacity.

Such a ‘pulsed inhibition’ (Klimesch et al., 2007; Mathewson et al., 2011) model of subitizing can be contrasted with an aperiodic account in which subitizing capacity is influenced by the overall excitation:inhibition (E:I) balance in the relevant brain areas. E:I balance is important for the stable maintenance of neural information, such as in working memory (Lim and Goldman, 2013; Barron et al., 2016), and may serve a similar function in object individuation. Recent work has shown through direct neural recordings, pharmacological manipulation, and computational modeling that the slope of the aperiodic 1/f^x^ power spectrum of electrophysiological brain activity provides an indirect measure of E:I balance (He et al., 2010; Gao et al., 2017; Lombardi et al., 2017). Thus, aperiodic slope provides an avenue to examine the relationship between E:I balance and subitizing capacity.

Here, we had participants undertake an enumeration task while we measured pre-stimulus neural activity with EEG. We characterised each individual’s subitizing range and related this to their pre-stimulus alpha peak frequency, power, and aperiodic slope. To anticipate the results, there was no relationship between alpha oscillations and subitizing capacity. By contrast, we observed a robust association between the individuals’ subitizing capacity and aperiodic slope, suggesting subitizing limits are influenced by the E:I balance of the visual system.

## Method

### Experimental Design

#### Participants

We aimed to collect at least fifty useable datasets, as this sample size yielded reasonable power to detect moderate correlations. Using G*Power 3 (Faul et al., 2007), we calculated that a sample size of 50 yielded 80% power to detect correlations of.375 and above using a two-tailed Spearman correlation. A recent meta-analysis of the relationship between alpha peak frequency and the temporal properties of vision estimated the true effect to be a correlation between *r* =.39 and *r* =.53 (95% bootstrap confidence interval; (Samaha and Romei, 2023), suggesting we were well powered to detect reasonable effect sizes for this effect. A total of 68 healthy participants were recruited from The University of Queensland’s online research participation platform. Data from seventeen participants were excluded from analysis. Five were excluded for not meeting our a priori criterion of at least 65% accuracy for enumerating a set-size of two items. Six were excluded for providing behavioural data that could not be fit adequately by the behavioural modelling procedure (described below), using a minimum criterion of R^2^ =.70, as suggested by Leibovich-Raveh and colleagues (Leibovich-Raveh et al., 2016). Finally, six participants were excluded from the EEG analyses as their neural data showed no discernible alpha oscillations (as determined by the specparam package, described below). All participants were compensated at a rate of $20 per hour for their time. Participants provided written informed consent prior to participation. All participants had normal or corrected-to-normal vision. The final sample contained 51 participants (37 female; mean age [SD] = 22.69 [3.87]; all right-handed). Demographic data were lost for two participants due to a technical error. The study was approved by the Human Research Ethics Committee at The University of Queensland.

#### Stimuli and Apparatus

The experiment was presented and controlled using Psychtoolbox-3.0.17 under MATLAB (R2020b; MathWorks), on a 22.5-inch VIEWPixx monitor (VPixx Technologies) with a 1920×1080 resolution and 100Hz refresh rate. Participants sat approximately 57cm away from the display, with a chinrest to help maintain this position. An Eyelink 1000Plus (SR Research, Ontario, Canada) video-based infra-red eye-tracker was used to ensure participants fixated centrally at the beginning of the trial. Participants responded using a USB keyboard.

Before each trial a red (RGB: 200, 0, 0) fixation cross (0.22° × 0.22°; thickness of 0.06°) appeared, centrally. During stimulus presentation (**Figure 1**), 16 coloured, vertical and horizontal bars (0.84° × 0.42°) appeared on a grey background (RGB: 160, 160, 160) for 30ms. Of these 16 bars, 2-8 were targets, with the remainder being distractors. Distractors were employed to ensure the amount of information presented was constant in all conditions. Bars were blue (RGB: 80, 80, 233) or green (RGB: 80, 142, 80) and of equal luminance (green = 21.03cd/m^2^, blue = 20.93cd/m^2^) as measured by a ColorCAL MKII colourimeter (Cambridge Research Systems). Orientation was varied between target and distractors because equiluminant colours are difficult to distinguish at short presentation durations. Colour-orientation pairing and target features were counterbalanced between participants, creating four possible target conditions (blue-vertical, blue-horizontal, green-vertical, and green-horizontal), with opposite colour-orientation pairing assigned to the distractors (i.e., green-horizontal, green-vertical, blue-horizontal, and blue-vertical, respectively). Target and distractor features were held constant throughout the experiment for each participant.

**Figure 1.**
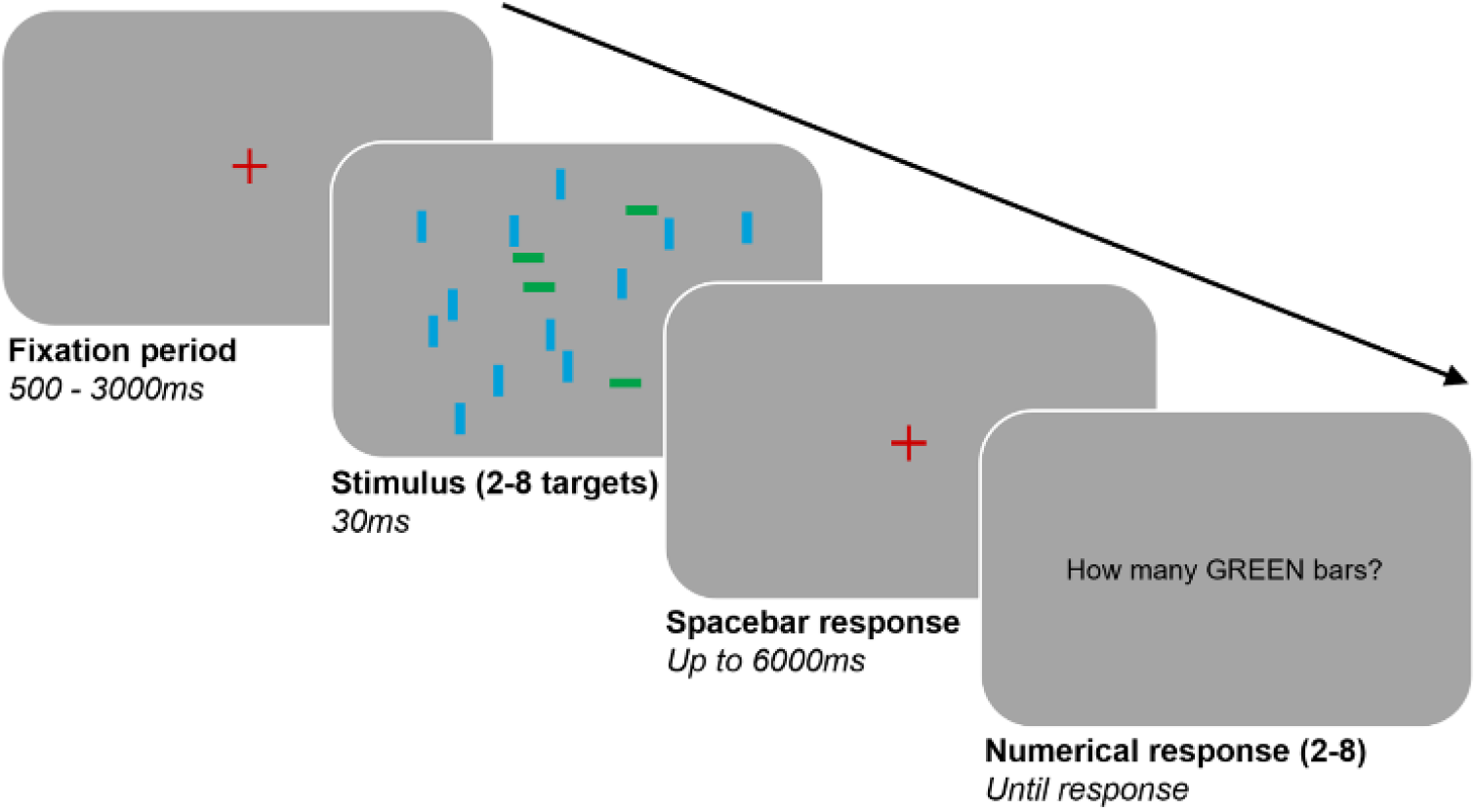
Stimuli and Trial Sequence. Example trial sequence. In the trial sequence shown here, the targets are green horizontal bars and the distractors are blue vertical bars. The number of target bars varied from 2-8, and there were always 16 bars in total. Size and placement of bars are not to scale. Participants were required to first press the spacebar when they had determined how many targets were present, then report the number of targets in the numerical response display.

Stimulus bars were placed within an imaginary annulus, 1.12° to 7.52° from the fixation cross, and spaced a minimum of 1.68° apart. The locations of the 16 stimulus bars were balanced across the display by allocating a randomly selected 4 bars to random locations within each quadrant of the display (subject to the above constraints).

#### Procedure

Participants completed the experiment in a dim room. Following EEG set-up (see below), the eye-tracker was calibrated using the standard 9-point calibration procedure.

After written and verbal instructions, participants completed one practice block, consisting of 28 trials. Practice trials were identical to experimental trials. There were 630 experimental trials, with 90 trials per set size, randomly ordered throughout the experiment. Breaks were provided after every 80 trials (roughly 5-7min), with the final block containing 70 trials.

Before each trial, participants had up to 3000ms to fixate on the red fixation cross for a minimum of 500ms. If fixation was unsuccessful, the eye-tracker was recalibrated. Once successful fixation was achieved, the stimuli were presented after a delay period of 1500-2500ms. Stimuli were displayed for 30ms. After 30ms, a red fixation cross replaced the stimulus display, and remained on screen until the participant pressed the spacebar to indicate they had enumerated the items for that trial. Their reaction time was taken as the time between stimulus onset and the press of the spacebar. This procedure was used to avoid the high RT variability that would come from employing a 7-alternative forced choice on the numerical responses. Participants were then prompted by written instructions on the display (e.g., “How many BLUE bars?”) to report the number of target bars using the keyboard number pad, and submitted their answer by pressing “Enter”. They were able to change their answer by re-selecting their preferred number before pressing “Enter”. Participants had up to 6000ms to press the spacebar before the trial was marked as an error, and the next trial was presented. No time limit was imposed during the numerical response. Experimenters monitored participants’ RTs and reinforced the instruction to finish counting before pressing the spacebar if participants routinely spent a long time selecting their numerical responses on larger set-size trials, indicating they may be continuing to count in this period. No accuracy feedback was provided.

#### EEG Acquisition

A 64-electrode BioSemi ActiveTwo system, digitised at 1024Hz with 24-bit A/D conversion, was used to record continuous EEG data. The 64 Ag/AgCl electrodes were arranged by the international standard 10-10 layout (Oostenveld and Praamstra, 2001) using a nylon head cap. All scalp electrodes were referenced to the Common Mode Sense and Driven Right Leg electrodes during recording, with the Driven Right Leg electrode as the ground electrode. Blinks were recorded with bipolar horizontal electro-oculographic (EOG) electrodes at the outer canthi of both eyes, and bipolar vertical EOG electrodes above and below the left eye, with bilateral mastoid electrodes as import references.

#### EEG Pre-Processing

The EEGLAB Toolbox (Delorme and Makeig, 2003) for MATLAB (R2020b) was used for offline EEG pre-processing. The data were down-sampled to 256Hz and high-pass filtered at 0.1Hz. With EEGLAB’s default Kurtosis-based electrode channel rejection, electrode channels with Kurtosis scores five SDs from the mean of all channels were rejected and interpolated. An average of 4% of electrodes were interpolated per participant. Data were re-referenced to the average of all electrodes, and epoched between -1000 to 2000ms relative to stimulus onset. The trial baseline (-100 to 0ms) was removed to aid the calculation of event-related potentials. The data were processed using an infomax Independent Component Analysis (ICA). The SASICA plugin for EEGLAB (Chaumon et al., 2015) was used to classify artifactual components such as blink artifacts, eye movements, muscle artifacts, and electrode noise. Trials containing blinks during stimulus presentation were removed from analysis (M[SD] = 3.8% [5.5%] of trials).

#### Electrode selection

To select electrodes for pre-stimulus analysis, we focussed on the electrode that showed the strongest post-stimulus response. To determine this, we first computed the Global Field Power (GFP; the standard deviation across electrodes at each sample throughout the trial) averaged across all trials and all participants, and found the time of greatest GFP. This time of greatest GFP was 172ms post stimulus. We then averaged the EEG data over a 100ms window centred on this time point and determined the electrode with the greatest absolute voltage over this window. This channel was PO8, with an average voltage of -3.76μV between 122 and 222ms post-stimulus (**Figure 2**). Thus, we used electrode PO8 as our electrode of interest (EOI). Additionally, we repeated our analyses across all electrodes, correcting for multiple comparisons with cluster-based permutation tests (cluster threshold:.05; minimum cluster size: 2 channels; number of permutations: 2000; neighbourhood definition: within 56mm, resulting in neighbourhoods of 3-8 channels; (Maris and Oostenveld, 2007), to determine the generality of the results beyond the EOI.

**Figure 2.**
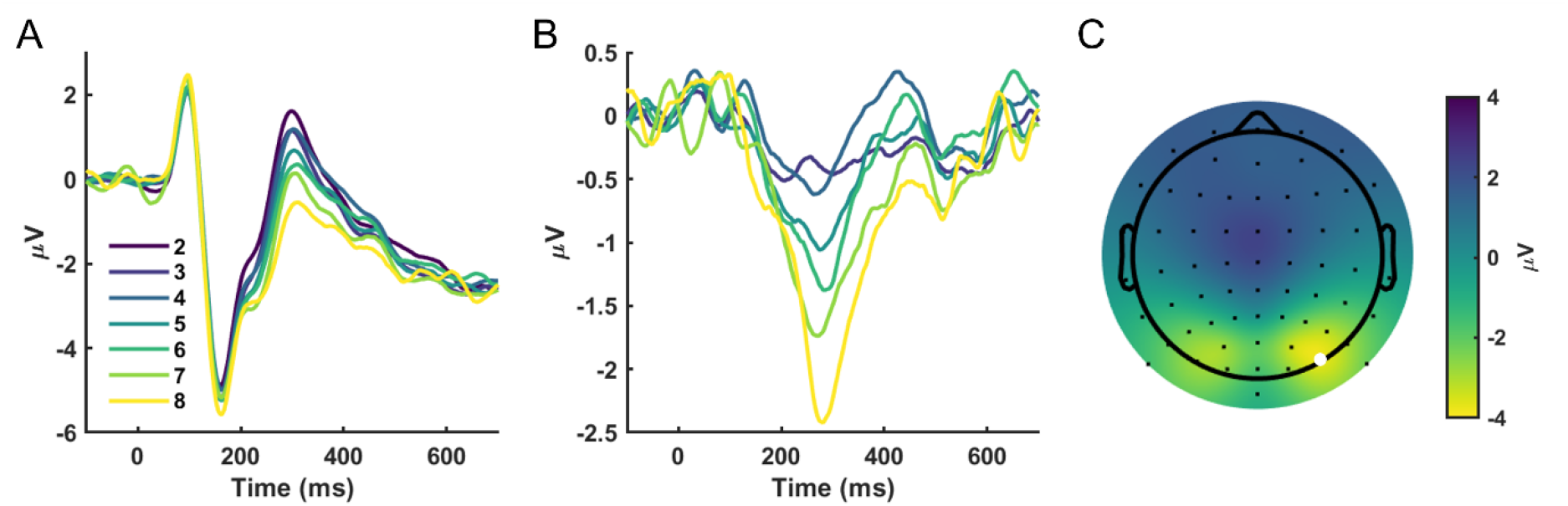
Event Related Potentials. **A**. ERPs at electrode PO8 for each set size. **B**. Difference waves from set-size 2. The ERP for set-size 2 has been subtracted from each ERP to illustrate the effect of increasing the number of target items on neural responses **C**. Average topography between 122-222ms, collapsed across set-size. The Electrode of Interest (PO8) is indicated as a white dot.

### Statistical Analyses

#### Behaviour

All analyses excluded trials with RTs greater than 2.5 SDs beyond the mean RT for each set size. On average, 2% (SD = 0.59) of trials were excluded per participant. Individual subitizing capacity, enumeration slope, and the intercept (reflecting the reaction time at the lowest set size) were calculated using a bilinear-sigmoid model (Leibovich-Raveh et al., 2016) on the RT (limited to correct trials) and accuracy data. First, a sigmoid function was fit to individual-level data, using a nonlinear error minimisation algorithm (the nlinfit function in MATLAB, employing the Levenberg-Marquardt algorithm). Then, two linear functions were fit: (1) a function with zero slope, at the sigmoid function’s y-intercept, and (2) a tangent line at the inflection point of the sigmoid function. Subitizing capacity was taken as the x-coordinate where the linear functions intersected. The slope of the enumeration function was extracted as the gradient of the tangent line. We refer to this as the ‘enumeration slope’ throughout, to distinguish it from the slope of the aperiodic function of the EEG (see below), which we refer to as ‘aperiodic slope.’

#### EEG

Pre-stimulus EEG data from -1000ms to stimulus onset at 0ms were used for all our analyses, to give an indication of the state of the neural system around the time of stimulus onset that was not confounded by stimulus-evoked brain responses. The data from this pre-stimulus period were Hann tapered and zero-padded to provide a frequency resolution of 0.25 Hz, and then used to compute power spectra for each trial with Welch’s method (Welch, 1967). For individual differences analyses, these spectra were averaged across all trials and analysed with specparam version 1.0.0 (Donoghue et al., 2020b) to extract periodic (oscillatory) and aperiodic spectral features. Specparam was run on data from 2-40 Hz with the following settings: peak width limits: 0.5-9 Hz; maximum number of peaks: 4; minimum peak height: 0 (which sets a peak height threshold relative to the scale of the data); peak threshold: 2; aperiodic mode: ‘fixed’. The model fits were good, yielding an average R^2^ of 0.993 (*SD* = 0.013). From these models, we examined the pre-stimulus individual alpha peak frequency, alpha power, and slope of the aperiodic function. Aperiodic offset was not analysed due to its extremely high correlation with aperiodic slope (*r* >.8), making its inclusion redundant. Spearman’s rank correlations and multiple regression analyses were used to examine the relationship between each of these neural parameters and the parameters of the enumeration function (i.e., the intercept, subitizing capacity, and the enumeration slope).

For within-participants analyses, specparam fits were calculated on the single-trial pre-stimulus EEG data and used to bin the trials into quartiles based on each of the neural parameters. Enumeration parameters were then computed from the trials in each of these bins, and the parameter estimates were fitted with linear mixed-effects models with fixed effects of quartile and random effects of participant to test for linear relationships between the neural parameters and enumeration outcomes.

## Results

### Behaviour

The reaction time results replicated the typical pattern for visual enumeration tasks (**Figure 3A**). Parameter estimates are presented in **Figure 3B**, with summary statistics shown in **Table 1**. The average estimated subitizing range was between 3 and 4 items, replicating the well-established capacity of subitizing (Jevons, 1871; Mandler and Shebo, 1982; Piazza et al., 2011). Error rates increased with set size and produced subitizing parameter estimates roughly in line with the RT results (Mean = 3.86; SD = 0.98). However, as our task was designed to emphasise reaction time over accuracy, we focused our analyses on correlating neural data with the parameters estimated from the reaction time results. Each of the enumeration parameters showed good reliability when parameters calculated from even- and odd-numbered trials were correlated (**Table 1**).

**Table 1.** Descriptive summary statistics (M, SD), reliabilities (r), and Spearman’s correlations between all variables, with neural measures taken from the EOI.

| Variables | M | SD | r | 1 | 2 | 3 | 4 | 5 |
| --- | --- | --- | --- | --- | --- | --- | --- | --- |
| **Reaction Time** |  |  |  |  |  |  |  |  |
| 1. Intercept | 0.79 | 0.14 | .93*** | - |  |  |  |  |
| 2. Subitizing Range | 3.54 | 0.41 | .80*** | .28*** | - |  |  |  |
| 3. Enumeration Slope | 0.29 | 0.05 | .78*** | -.03 | .45*** | - |  |  |
| **Neural** |  |  |  |  |  |  |  |  |
| 4. Alpha Peak Frequency | 11.14 | 0.99 | .96*** | .04 | -.16 | -.07 | - |  |
| 5. Alpha Power | 0.77 | 0.38 | .99*** | .15 | -.03 | .21 | -.20 | - |
| 6. Aperiodic Slope | 1.51 | 0.26 | .99*** | -.06 | .34** | .30* | -.39** | .40** |

Note: \* $p < .05$ , \*\* $p < .01$ , \*\*\* $p < .001$

**Figure 3.**
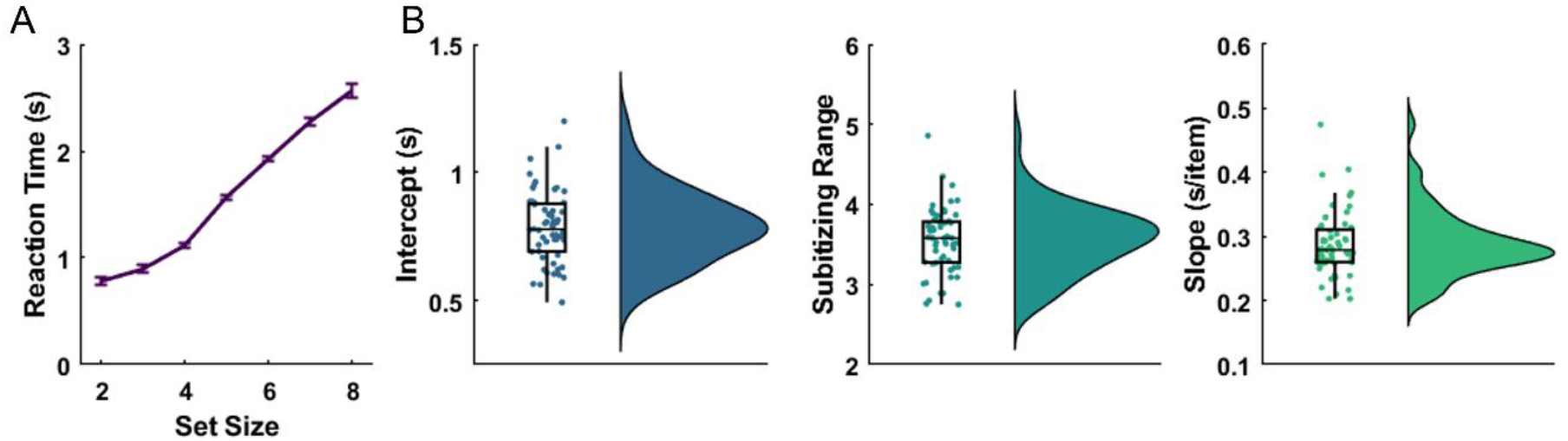
Behavioural Results. **A**. Reaction time results. **B**. Box plots and densities representing the enumeration model outputs for each individual (small dots), fit to the reaction time data. Boxes represent the interquartile range, and the central line reflects the median. Whiskers reflect 1.5 times the interquartile range, or the most extreme value, whichever is smaller. Error bars in A represent within-participants standard error of the mean. Plots created with code adapted from (Allen et al., 2019).

### Individual differences

Each of the neural parameters showed good reliability when parameters calculated from even- and odd-numbered trials were correlated (**Table 1**). To examine the relationship between enumeration properties, in particular the individual subitizing range, and pre-stimulus neural state properties, we first performed Spearman’s rank correlations between each pair of variables, with neural measures taken from the EOI (**Table 1**). This revealed significant correlations between aperiodic slope and both alpha power and peak frequency. Aperiodic slope was also significantly positively correlated with enumeration slope and the individual subitizing range. Finally, subitizing range was correlated with both enumeration slope and the intercept of the enumeration function. Estimation of subitizing parameters with an alternative method (bilinear fit) yielded the same results.

We next included each of the neural parameters in a multiple regression analysis to predict enumeration performance. In our multiple regression analyses, each parameter of the enumeration model served as the criterion in distinct analyses, with the three neural state measures as predictors. To control for any deviations from normality, we performed these regressions on the ranked data, as in the Spearman’s rank correlations above.

To confirm that the relationships between aperiodic slope and alpha oscillation measures did not produce problematic multicollinearity, we assessed them using the variance inflation factor (VIF). The VIF has a minimum of 1, and values above 5 are typically interpreted as indicating a confounding influence in regression analyses (Akinwande et al., 2015). Here, the VIF values produced for each of our predictors in the context of the others were well below 2 (alpha peak frequency = 1.21, alpha power = 1.16, aperiodic slope = 1.33), indicating their suitability for inclusion in the multiple regression.

The multiple regressions produced non-significant models for the intercept (baseline reaction time) and the slope of the enumeration model (*p*s >.33). For subitizing range, a significant model was produced, R^2^ =.22, *p* =.009, with the sole significant predictor being the direct effect of aperiodic slope on the individual subitizing range, beta = 0.44, *p* =.004 (**Figure 4**), such that increased subitizing range was predicted by an increased aperiodic slope (i.e., increased inhibition). Exploration of more complex models including all the possible two- and three-way interactions between the predictors produced the same results: the models predicting intercept and enumeration slope were nonsignificant (*p*s >.31), but the model predicting individual subitizing range was significant (R^2^ =.36, *p* =.005), with aperiodic slope being the only significant predictor (beta = 1.38, *p* =.025). There was little difference in model selection criteria between the two subitizing range models: AIC was slightly higher for the simpler model (Additive: 420.73; Interactive: 418.56) but BIC was somewhat lower (Additive: 428.46; Interactive: 434.01).

**Figure 4.**
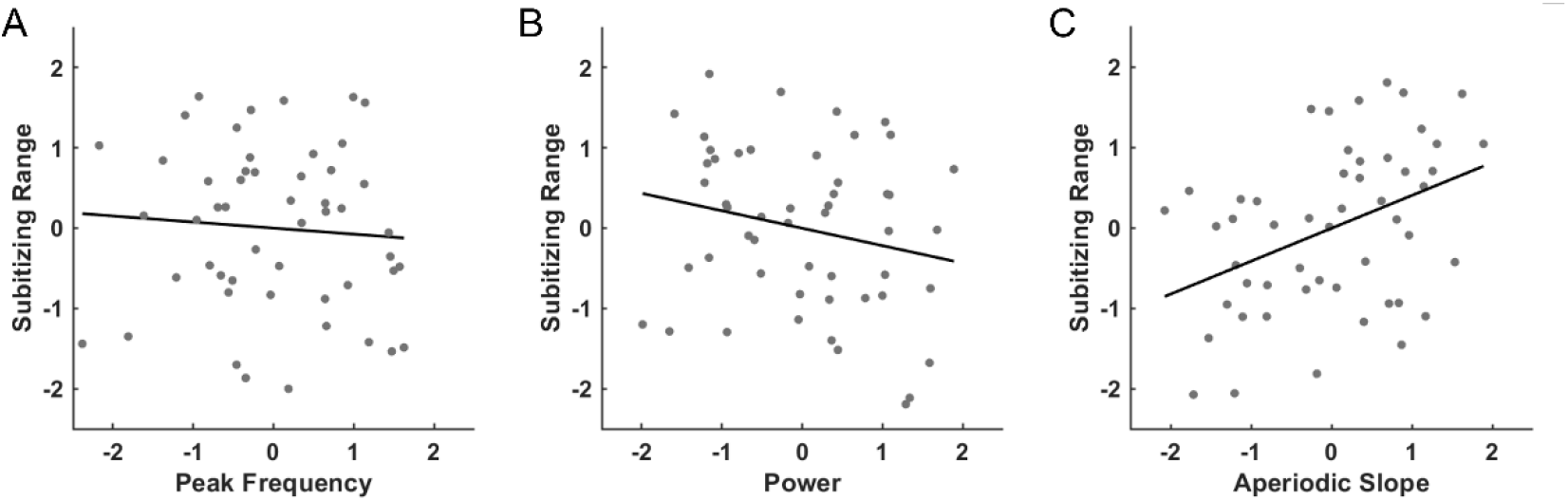
Standardised relationships between subitizing range and each predictor, controlling for all other predictors. Shown are the relationships between Individual Subitizing Range and **A**. Alpha Peak Frequency, **B**. Alpha Power, and **C**. Aperiodic Slope. Each plot displays the partial correlation between its predictor on the x axis and Subitizing Range on the y axis, after controlling for the other two predictors.

To explore the spatial distribution of these effects outside the EOI we repeated the multiple regression analysis at every electrode, correcting for multiple comparisons with cluster permutation. To simplify the interpretation of results, we used the additive model with no interactions for this analysis. The analysis of neural predictors of subitizing range produced one significant cluster (*p*_cluster_ =.006), made up of 3 right-posterior electrodes (O2, PO4, and PO8). Examining the coefficients at these electrodes, none showed significant relationships between subitizing range and alpha peak frequency. One electrode showed a significant effect of alpha power (*p* =.02), and two showed significant effects of aperiodic slope (*p*s <.011). There were no significant clusters relating the neural measures to the intercept or slope of the enumeration function.

### Within Participants Effects

To assess the relationships between the parameters of the enumeration function and each of our neural predictors within participants, we computed alpha peak frequency, alpha power and aperiodic slope in the pre-stimulus period of single trials. To quantify within-participants relationships, we divided single-trial measures of each of these variables into quartiles and fit the behavioural data from each quartile with the linear-sigmoid enumeration function. This yielded estimates of the intercept, subitizing range and enumeration slope for each quartile within each participant. These parameter estimates served as the criterion values to be predicted by a linear mixed effects model with a fixed effect of quartile label and random effect of participant. This allowed us to quantify consistent linear relationships between neural predictors and enumeration parameters. The results are shown in **Table 2**.

**Table 2.** Within Participants results for each enumeration parameter being predicted by each neural measure.

| Predictor | Intercept |  | Subitizing Range |  | Enumeration Slope |  |
| --- | --- | --- | --- | --- | --- | --- |
| | $\beta$ | $p$ | $\beta$ | $p$ | $\beta$ | $p$ |
| Alpha Peak Freq | 0.04 | 0.011* | 0.05 | 0.728 | 0.22 | 0.059 |
| Alpha Power | -0.03 | 0.101 | 0.23 | 0.108 | -0.02 | 0.681 |
| Aperiodic Slope | -0.04 | 0.002** | -0.06 | 0.544 | 0.01 | 0.794 |

Note: \* $p < .05$ , \*\* $p < .01$ , \*\*\* $p < .001$

There was a significant negative relationship between aperiodic slope and the intercept of the enumeration function, indicating that as aperiodic slope increased reaction times were faster at the lowest set-size (**Figure 5**). There was also a significant positive association between alpha peak frequency and intercept, indicating that trials with higher alpha peak frequency (i.e., faster alpha oscillations) were associated with faster reaction times at the smallest set size. No associations were observed between the neural parameters and individual subitizing range or enumeration slope.

**Figure 5.**
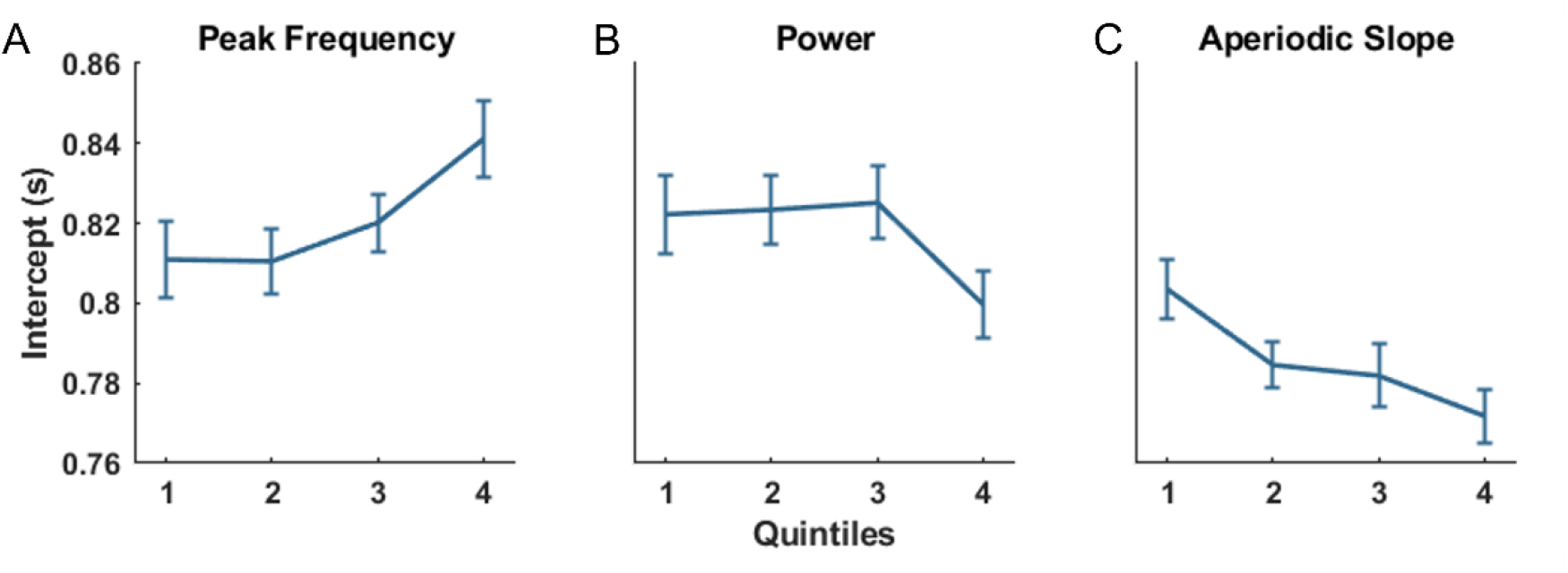
Within participants relationship between the intercept of the enumeration function and each of the neural predictors. Shown are the relationships between Individual Subitizing Range and **A**. Alpha Peak Frequency, **B**. Alpha Power, and **C**. Aperiodic Slope. Each of the neural predictors was divided into quintiles and the enumeration model was fit to the behavioural data from the trials in each quintile. Only the intercept of the enumeration function produced significant linear associations with the neural predictors (see Table 2). Error bars represent within-participants standard error of the mean.

Repeating this analysis at every electrode and correcting for multiple comparisons with cluster permutation revealed no significant relationships between the neural parameters and either the intercept or subitizing range (although subitizing range showed a marginal positive relationship with aperiodic slope, *p*_cluster_ =.067). Enumeration slope was significantly negatively related to alpha peak frequency at a cluster of two central electrodes (Cz & CP1; *p*_cluster_ =.001), indicating that as alpha oscillations slowed and cycle durations increased, responses became progressively slower at higher set sizes. Enumeration slope was also significantly positively related to aperiodic slope at a cluster of three central electrodes (FC1, Cz & CP2; *p*_cluster_ =.040), indicating reaction times were slowed more at higher set sizes when aperiodic slope (i.e., inhibition) was high. No other relationships with enumeration slope were significant.

## Discussion

We set out to test whether alpha oscillations underlie capacity limits in visual object individuation, and contrasted this with an account relating individuation to E:I balance assessed via aperiodic 1/f^x^ slope. The results were clearly in favour of the E:I balance account. Individual differences in subitizing capacity were associated with pre-stimulus aperiodic slope but not pre-stimulus alpha peak frequency. The within-participants analysis showed a trend towards the same result. These results were observed in parieto-occipital channels, consistent with the previously reported parietal locus of object individuation (Xu and Chun, 2009). Both pre-stimulus aperiodic slope and alpha peak frequency were associated within participants with changes in the intercept of the enumeration function, which reflects the reaction time in the easiest, lowest set-size trials. These results are consistent with previous findings that alpha oscillations correspond to changes in subjective perceptual thresholds, rather than the quality or efficiency of visual information processing (Samaha et al., 2020). Alpha peak frequency was also related to the slope of the enumeration function outside the subitizing range at more central electrode sites. Together, these results speak against alpha-frequency sampling as the source of perceptual capacity limitations and instead support aperiodic activity as an important indicator of visual processing influences such as E:I balance.

A steeper aperiodic slope is an indication of skew in E:I balance toward increased inhibition (Gao et al., 2017; Molina et al., 2020; Ahmad et al., 2022; Wiest et al., 2023; Salvatore et al., 2024). Our observation of a positive association between aperiodic slope and subitizing capacity suggests that more items can be subitized with greater levels of inhibition in the cortical areas involved in the processing (presumably within healthy limits). There are a number of computational mechanisms that might explain such a result. Small increases in inhibition can lead to noise reduction in neural systems, enhancing the signal to noise ratio of task-related signals (Tetzlaff et al., 2012). Noise suppression might allow more items to be processed before the neural signal is degraded by neural activity that is unrelated or stochastic. Such a framework would be consistent with previous findings that suggest rapid sequential processing even within the subitizing range (e.g., Wutz et al., 2012; Wutz and Melcher, 2013). Increased neural inhibition is also associated with more efficient distractor suppression (Zhang et al., 2014). This is unlikely to explain our individual differences results because distractor suppression should enhance target processing generally, rather than influencing the subitizing range specifically. Distractor suppression might explain the within-participants results relating aperiodic slope to the intercept and slope of the enumeration function, however, as we discuss below. Finally, inhibition-related aperiodic neural activity may have been associated with subitizing capacity via its role in attentional selection. Attention and the subitizing range are robustly related (Chen et al., 2022), and there is emerging evidence that aperiodic activity is a robust correlate of attentional states (Waschke et al., 2021; Pietrelli et al., 2022). It may be that items outside the current focus of attention, both targets and distractors, are suppressed by aperiodic activity, having their processing delayed in such a way that reaction times increase dramatically beyond attentional capacity (Reynolds and Chelazzi, 2004). Interestingly, we have shown in recent work that the perceptual consequences of lateral inhibition are positively associated with aperiodic slope (Williams et al., 2024).

Interestingly, pre-stimulus aperiodic activity was associated with both the intercept and slope of the enumeration function within individuals. We made no explicit predictions about these parameters, so these findings should be taken as exploratory observations; however, we can speculate about their implications. The intercept relationship was over the electrode of interest (PO8) that was selected to capture early elements of the enumeration process. The relationship to enumeration slope, however, was observed over more central electrode sites. This dissociation of effect location implies a difference of source and possibly process. The central location suggests a possible involvement of attentional control in counting or estimating higher set-size displays. We might expect that as attention is required to shift around the mental representation of the stimulus array, to process items that could not be individuated within the first sweep of visual processing, fronto-central regions involved in attentional control become more involved in the task. Pre-stimulus aperiodic slope was negatively associated with the enumeration intercept, suggesting that at the lowest set-sizes, participants were able to respond faster when there was more inhibition in the system. However, performance outside the subitizing range was slowed by increased inhibition prior to stimulus onset, as demonstrated by the positive relationship between aperiodic and enumeration slopes. These results demonstrate the complex, ‘more does not mean better,’ relationship between neural inhibition and task performance (for a similar interpretation of alpha-related inhibition, see also (Lange et al., 2013).

While our results suggest that alpha frequency sampling is not the source of perceptual capacity limits, these results do not argue against the involvement of alpha oscillations in other elements of enumeration or other related tasks. We showed within-participants relationships between alpha peak frequency and both the intercept and slope of the enumeration function, clearly indicating alpha’s involvement in other aspects of task performance. Alpha’s association with these two distinct parameters may reflect distinct roles played by alpha oscillations in this task. Alpha’s association with the intercept parameter, which indicates participants fastest reaction times, may stem from alpha’s clear association with individual response thresholds in visual tasks (Limbach and Corballis, 2016; Iemi et al., 2017). The observed link between alpha and the slope of the enumeration function, however, is more likely to be due to alpha’s role in selective spatial attention (Klimesch, 2012; Peylo et al., 2021). Such links to attention would be consistent with previous findings of an association between alpha burst-rate and task load up to perceptual capacity in multi-object tracking (MOT) tasks (Wutz et al., 2020). MOT performance reflects a similar perceptual capacity to that assessed with enumeration paradigms (Eayrs and Lavie, 2018). It is unclear why work with MOT has shown relationships between tracking capacity and alpha oscillations, whereas the current work does not. One possibility is that, due to its dynamic nature, MOT may produce the appearance of a hard boundary on neural activity related to task performance, whether or not that activity is involved in producing the perceptual capacity limit. In enumeration, participants can continue to count or estimate items beyond their perceptual capacity, albeit with impaired performance. In MOT, items beyond this limit must be abandoned to enable the participant to track the within-capacity items. Thus, neural measures related to task performance are likely to plateau at perceptual capacity limits in MOT tasks (e.g., Drew and Vogel, 2008; Alnæs et al., 2014), whether they are the source of such a limit or not. In the enumeration paradigm, however, this is not the case.

Importantly, the analysis approach we employed here divides oscillations and aperiodic patterns into dissociable components of the neural signal (Donoghue et al., 2020b). It has recently been established that failure to separate these elements can result in biased estimation of both alpha power and peak frequency (Donoghue et al., 2020a; Samaha and Cohen, 2022). It may be noted that, despite this separation, aperiodic slope was still moderately correlated with the alpha parameters. This is unsurprising, both methodologically and conceptually. While our fit statistics showed our models had an excellent fit to the data, the parameter separation may not have been perfect. Past work has shown, however, using these same analysis methods, that separating aperiodic and periodic parameters reduces the bias in estimates of oscillatory power and peak frequency (Donoghue et al., 2020a; Samaha and Cohen, 2022), so it is unlikely these correlations were introduced by our analysis. Instead, it is possible that the correlation between alpha oscillations and aperiodic slope is veridical. Alpha oscillations are well established as reflecting inhibitory processes in visual processing (Klimesch et al., 2007; Händel et al., 2011; Lange et al., 2013; Borghini et al., 2018). As aperiodic slope is an indirect measure of the E:I balance in the system, it likely encompasses some inhibitory effects related to alpha oscillations, but occurring outside the alpha frequency range, such as tonic effects, and cross-frequency interactions (Canolty and Knight, 2010; Voytek et al., 2010). It is unlikely these relationships affected the results we report here. Our regression analyses produced results that uniquely reflect the effect of aperiodic slope. Additionally, analysis of the Variance Inflation Factor confirmed that the pre-stimulus neural measures influenced each other minimally in the regression that formed the basis of our individual differences analysis.

In summary, perceptual capacity limits do not seem to arise from temporal constraints on alpha-frequency sampling of visual information. Rather, they are associated with non-rhythmic aspects of processing that bias the E:I balance of the neural system toward inhibition. Both alpha oscillations and aperiodic slope show robust relationships with inhibition in neural systems, that nonetheless dissociate in this task. This dissociation may be due to associations with different inhibitory mechanisms or neuron types, or it may reflect the different temporal properties of the inhibition involved.

## Acknowledgements

The authors would like to acknowledge the following funding sources: AMH was supported by the Australian Research Council (DE220101019). JBM was supported by a National Health and Medical Research Council (Australia) Investigator Grant (2010141). The authors would also like to thank paper-wizard.com for helpful comments on an earlier version of this manuscript.

